# Perceptual uncertainty explains activation differences between audiovisual congruent speech and McGurk stimuli

**DOI:** 10.1101/2023.09.10.556693

**Authors:** Chenjie Dong, Uta Noppeney, Suiping Wang

**Author notes:** Senior authors contributed equally to this work and are listed in alphabetical order (Uta Noppeney, Suiping Wang). **Correspondence should be addressed to**: Suiping Wang. **Author contributions** Chenjie Dong, conceptualization, data collection, data analysis, writing manuscript; Uta Noppeney, conceptualization, resources, writing manuscript, supervision; Suiping Wang, conceptualization, resources, writing manuscript, supervision, funding acquisition, project administration.

## Abstract

Face-to-face communication relies on the integration of acoustic speech signals with the corresponding facial articulations. While the McGurk illusion is widely used as an index of audiovisual speech integration, critics argue that it arises from perceptual processes that differ categorically from natural speech recognition. Conversely, Bayesian theoretical frameworks suggest that both the illusory McGurk and the veridical audiovisual congruent speech percepts result from probabilistic inference based on noisy sensory signals. According to these models, the inter-sensory conflict in McGurk stimuli may only increase observers’ perceptual uncertainty. This functional magnetic resonance imaging (fMRI) study presented participants (20 male and 24 female) with audiovisual congruent, incongruent, and McGurk stimuli along with their unisensory counterparts in a syllable categorization task. Behaviorally, observers’ response entropy was greater for McGurk compared to congruent audiovisual stimuli. At the neural level, McGurk stimuli increased activations in a widespread neural system, extending from the inferior frontal sulci (IFS) to the pre-supplementary motor area (pre-SMA) and insulae, typically involved in cognitive control processes. Crucially, in line with Bayesian theories these activation increases were fully accounted for by observers’ perceptual uncertainty as measured by their response entropy. Our findings suggest that McGurk and congruent speech processing rely on shared neural mechanisms, thereby supporting the McGurk illusion as a valid measure of natural audiovisual speech perception.

**Significance Statement:** Effective face-to-face communication relies on integrating acoustic speech signals with the corresponding facial articulations. While McGurk illusion is extensively used to study audiovisual speech perception, recent critiques argue that it may be categorically different from typical speech recognition because of the conflict between the audiovisual inputs. This study demonstrates that McGurk stimuli increase activations in a network of regions typically involved in cognitive control. Crucially, the activation differences between McGurk and normal speech stimuli could be fully accounted for by the variation in observers’ perceptual uncertainties. Our results suggest that McGurk and congruent audiovisual speech stimuli rely on shared neural mechanisms – thereby supporting the validity of the McGurk illusion as a tool for studying natural audiovisual speech perception.

## Introduction

Effective face-to-face communication relies on the integration of auditory speech with the corresponding facial articulations. In laboratory settings, the McGurk illusion is often employed to study audiovisual integration of speech (McGurk and MacDonald, 1976). This illusion arises when a discrepancy is introduced between the visual and auditory speech signals. For instance, when an auditory ba phoneme is presented simultaneously with a facial articulation of a ga (i.e., viseme), observers frequently perceive an illusory auditory ‘da’ percept.

Despite its widespread use, the relevance of the McGurk illusion for understanding the mechanisms of natural audiovisual speech comprehension has recently been questioned (Getz and Toscano, 2021; Van Engen et al., 2022). Critics argue that McGurk stimuli categorically differ from natural speech stimuli because they introduce a conflict between auditory and visual signals (Erickson et al., 2014; Getz and Toscano, 2021; Van Engen et al., 2022) – thereby invoking additional conflict monitoring and cognitive control processes. In line with this view, neuroimaging studies showed that McGurk stimuli activated not only posterior superior temporal gyri and sulci (pSTG/S) (Bernstein et al., 2008; Benoit et al., 2010; Baum et al., 2012; Szycik et al., 2012; Luttke et al., 2016), areas traditionally associated with audiovisual speech processing, but also anterior cingulate cortex (ACC) /pre-supplementary motor area (pre-SMA) (Moris Fernandez et al., 2017; Moris Fernandez et al., 2018; Murakami et al., 2018), and inferior frontal gyri/ sulci (IFG/S) (Hasson et al., 2007; Skipper et al., 2007; Tse et al., 2015; Gau and Noppeney, 2016; Murakami et al., 2018), i.e. regions implicated in conflict monitoring and cognitive control.

In contrast, Bayesian theoretical frameworks propose that veridical percepts for audiovisual congruent stimuli and illusory percepts for McGurk stimuli emerge from common computational mechanisms (Noppeney and Lee, 2018; Noppeney, 2021; Shams and Beierholm, 2022). In both instances, observers need to infer the phoneme-viseme pair and its underlying causal structure from audiovisual signals that are corrupted by noise (Magnotti and Beauchamp, 2015, 2017). Various internal and external noise sources may thus introduce discrepancies between audiovisual inputs on congruent trials, while on McGurk trials they may eliminate discrepancies. Thus, the noisy inputs for McGurk and congruent stimuli may even be identical on a particular trial, even though the true underlying audiovisual phoneme-viseme pairs differ. Indeed, recent research shows that when observers perceive an illusory ‘da’ percept on McGurk trials, they often incorrectly infer a congruent relationship between these physically conflicting audiovisual signals. Conversely, they can incorrectly infer incongruent relationships between a phoneme and a viseme, when they miscategorized the phoneme on congruent trials (Meijer and Noppeney, 2023).

These findings highlight the inherent uncertainty observers face when making perceptual decisions about phonemes, visemes and their causal relationship on both congruent and McGurk trials. For both congruent and McGurk stimuli, perceptual inference relies on the integration of noisy multisensory information with prior knowledge. Yet, despite the shared computational mechanisms, the perceptual decisions on McGurk trials may on average be associated with greater uncertainty compared to trials on which audiovisual information is congruent (Kimmet et al., 2023; Meijer and Noppeney, 2023). Based on this theoretical framework we hypothesized that the increased SMA and IFS activations during McGurk trials may be explained by observers’ greater perceptual uncertainty when presented with conflicting than congruent audiovisual signals.

To test this hypothesis, this fMRI study presented observers with audiovisual congruent (AVc), incongruent (AVi), and McGurk (AVm) phoneme-viseme pairs along with their unisensory counterparts in a syllable categorization task. First, we assessed activation differences for McGurk stimuli compared to congruent and incongruent phoneme-viseme pairs. Second, we examined whether BOLD responses varied for different syllables within each sensory context (i.e., auditory, visual, audiovisual congruent). Third, we investigated whether variations in observers’ perceptual uncertainty, as reflected in the entropy of their response distributions, can account for the activation differences between McGurk and congruent/incongruent pairs as well as across different syllables.

## Materials and Methods

### Participants

Forty-four healthy participants (24 women, mean age = 20.86, SD = 2.27) were recruited from South China Normal University. Two participants (2 women) were excluded from the data analysis as they were unable to follow the experimental instruction. All participants had normal or corrected-to-normal vision, no history of neurological or psychiatric disorders, and provided informed written consent before the experiment. The protocol for this study was approved by the Human Research Ethics Committee of the School of Psychology at South China Normal University.

### Materials

Short clips recorded from a male actor were presented to the participants. The clips included three auditory stimuli (A: A_ba_, A_da_, A_ga_), three visual stimuli (V: V_ba_, V_da_, V_ga_), three audiovisual congruent stimuli (AVc: AVc_ba_, AVc_da_, AVc_ga_), one McGurk stimulus (AVm = A_ba_ + V_ga_), and one audiovisual incongruent stimulus (AVi = A_ga_ + V_ba_). The auditory stimuli were generated by removing the video of the audiovisual congruent stimuli; the visual stimuli were generated by removing the audio of the audiovisual congruent stimuli; the AVm stimuli were generated by dubbing an auditory ba over the video of facial articulations of ga; the AVi stimuli were generated by dubbing an auditory ga over the video of facial articulations of ba in Adobe Premiere.

The video clips had a duration of 2 seconds, were recorded at a frame rate of 25 frames per second and had a resolution of 640 × 480 pixels. The audio stimuli were recorded at a sampling rate of 48 kHz and were presented to the participants at an approximately sound level of 70 dB using MRI-compatible earphones. To ensure the intelligibility of the stimuli inside the MRI scanner, participants (n=28) completed a 6 - scale questionnaire about the intelligibility of the stimuli (1 - very unclear, 3 - clear, 6 - very clear) after scanning. The mean intelligibility of auditory stimuli was 5.15 (SD = 0.87), and the mean intelligibility of visual stimuli was 5.58 (SD = 0.61).

### Experimental design and procedures

In the syllable categorization task, participants were presented with auditory, visual, audiovisual congruent, incongruent, or McGurk stimuli and indicated the syllable they heard (or saw on visual trials) by pressing one of the 4 buttons (‘ba’, ‘da’, ‘ga’, and ‘others’) with index and middle fingers of their left and right hand (**figure 1**). The response fingers were counterbalanced both within and between participants. Each trial started with a fixation cross (0.5 s), followed by the A, V or AV stimulus (2 s), a blank screen (1.5 s), the four-alternative forced choice (4AFC) screen (2 s) and an inter-trial interval randomly sampled from 2 seconds, 4 seconds, or 6 seconds. A baseline condition (a white fixation on center of the screen (14 s) was presented at the beginning and end of each run. The A and V stimuli were presented in separate unisensory runs in counterbalanced order; the AV stimuli were presented in a randomized order in each audiovisual run. Each unisensory run included 30 trials (i.e., 10 repetitions per syllable, run duration = 8 minutes); each of the 4 audiovisual runs included 30 congruent trials (i.e., 10 repetitions per syllable), 20 McGurk trails, and 10 incongruent trials (run duration = 12 minutes).

**Figure 1.**
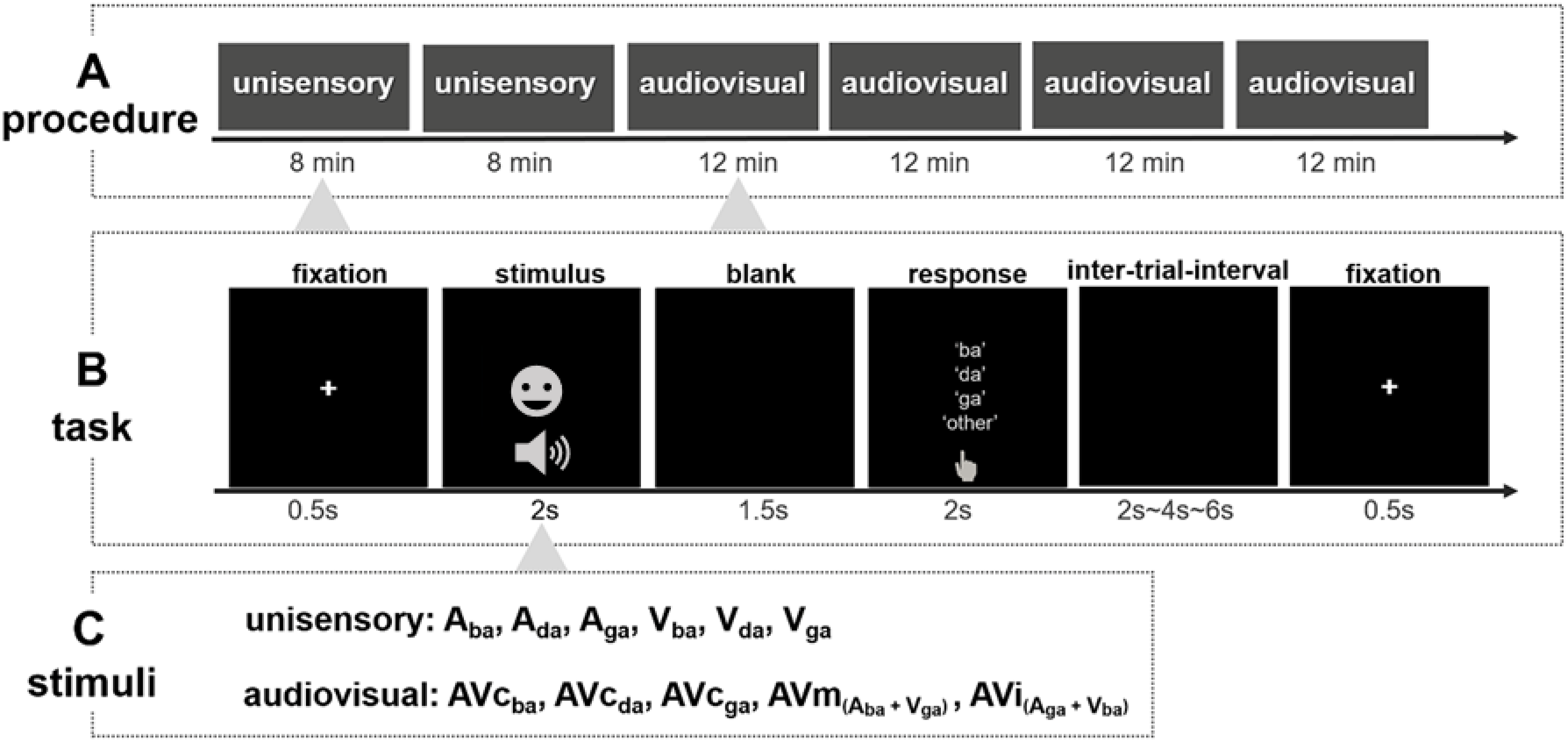
Experimental procedure and stimuli. (A) Experimental design. Participants were presented with auditory, visual, and audiovisual syllables. (B) Example trial. Participants were presented with an audiovisual movie (2 s) followed by a blank and response period. In a four alternative forced choice task they categorized the syllable as ‘ba’, ‘da’, ‘ga’, or ‘other’ (C) Set of unisensory and audiovisual stimuli. Abbreviations: A_ba_ = auditory ba, A_da_ = auditory da, A_ga_ = auditory ga, V_ba_ = visual ba, V_da_ = visual da, V_ga_ = visual ga, AVc_ba_ = audiovisual congruent ba, AVc_da_ = audiovisual congruent da, AVc_ga_ = audiovisual congruent ga, AVm = McGurk (i.e., auditory ba with visual ga), and AVi = audiovisual incongruent (i.e., auditory ga with visual ba).

### Behavioral data analyses

Behavioral data were analyzed using the JASP toolbox (https://jasp-stats.org) based on R. For each stimulus, we calculated the syllable categorization accuracy and Shannon entropy over the response distribution.

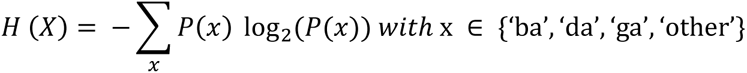

The Shannon entropy is maximal when observers randomly choose each of the four response options with equal probability (i.e., 25% for ‘ba’, ‘da’, ‘ga’, and ‘other’). While entropy is computed over trials, it is related to participants’ uncertainty on a particular trial. Hence, in this study we use response entropy as an index for observers’ perceptual uncertainty associated with different types of stimuli.

#### Categorization accuracy

To assess whether observers benefitted from audiovisual integration we entered categorization accuracy into a 3 (modality: A, V, and AVc) × 3 (syllable: ba, da, and ga) repeated measure ANOVA. Moreover, we assessed whether accuracies on unisensory and audiovisual congruent da stimuli are correlated with observers’ McGurk illusion rate using a Pearson correlation.

#### Entropy

We examined whether the audiovisual congruent stimuli reduced the perceptual uncertainty compared to the unisensory stimuli using a 3 (modality: A, V, and AVc) × 3 (syllable: ba, da, and ga) repeated measures ANOVA on entropy. Moreover, we assessed whether the McGurk stimulus was associated with greater uncertainty than the audiovisual congruent da stimulus by comparing the entropies of those two conditions in a paired t-test.

### MRI data acquisition

MRI data were collected at the Brain Imaging Center at South China Normal University on a Siemens 3T Prisma fit scanner with a 20-channel head coil. High-resolution T1-weighted anatomical images were collected using a multi-echo MPRAGE pulse sequence (repetition time [TR] = 2.53 s; echo time [TE] = 1.94ms, flip angle = 7°, field of view [FOV] = 256mm, matrix = 256 × 256, slice thickness = 0.5mm, slices number = 176). Functional data were collected using a T2*-weighted echo planar imaging EPI pulse sequence sensitive to blood oxygen level-dependent (BOLD) contrast (TR = 2s, TE = 30ms, flip angle = 90, FOV = 192 × 192mm, matrix = 64 × 64mm, slice thickness = 2mm, slice number = 62).

### MRI data analysis

#### fMRI data preprocessing

Data were analyzed using statistical parametric mapping (SPM12, Wellcome Department of Imaging Neuroscience, University College London; http://www. fil.ion.ucl.ac.uk/spm) (Friston et al., 1994) running on MATLAB 2021b. Scans from each participant were realigned using the first as a reference, spatially normalized into MNI standard space using parameters from segmentation of the T1 structural image (Ashburner and Friston, 2005), resampled to 2*2*2 mm^3^ voxels, and spatially smoothed with a Gaussian kernel of 6 mm FWHM. The time-series in each voxel was high-pass filtered to 1/128 Hz.

#### General linear model analysis

In all general linear models, data were modeled in an event-related fashion with regressors entering the design matrix after convolving each event-related boxcar (representing a single trial, duration of trial = 2s) with a canonical hemodynamic response function (HRF). Realignment parameters were included as nuisance covariates to account for residual motion artifacts.

Overall, we generated four general linear models: GLM-1A investigated the effects of sensory context by comparing A, V, AVc, AVm, and AVi (pooling over syllable categories). GLM-2A assessed the effect of syllable categories separately for A, V and AV sensory modalities. Finally, GLM-1B and GLM-2B investigated whether activation differences across sensory contexts or syllables can be explained away by differences in perceptual uncertainty as measured by Shannon entropy for each stimulus.

GLM-1A included six regressors (fixation baseline, A, V, AVc, AVm, and AVi). At the first (i.e. within subject) level, we computed the following contrasts. (1) A > baseline, V >baseline, (2) AVm > AVc, AVm > AVi, (3) AVm < AVc, AVm < AVi. The contrast images were entered into one sample t-tests at the 2^nd^ i.e., group level.

GLM – 2A included twelve regressors (fixation baseline, A_ba_, A_da_, A_ga_, V_ba_, V_da_, V_ga,_ AVc_ba_, AVc_da_, AVc_ga,_ AVm and AVi). At the first level, we computed contrasts for each condition relative to baseline. We entered the nine contrast images for three syllables (ba, da, ga) × (A, V, and AVc) into a second level repeated-measures ANOVA. In this repeated measures ANOVA, we assessed differences across the three syllables by computing F-tests across syllables separately for the auditory (A_ba_ vs A_da_ vs A_ga_), visual (V_ba_ vs V_da_ vs V_ga_), and audiovisual congruent (AVc_ba_ vs AVc_da_ vs AVc_ga_) stimuli.

GLM-3 includes only 2 regressors, one regressor modelling the onsets of all stimuli irrespective of sensory modality or syllable categories and a second regressor, a parametric modulator, that encodes Shannon entropy associated with each stimulus on each trial. The parameter estimates of the parametric modulator were entered into a 2^nd^ level one sample t-test to test for the effect of entropy.

GLM-1B and GLM-2B were equivalent to GLM1A and 2A, except that they additionally included a single parametric modulator that modelled the stimulus-specific entropy for each trial. We replicated the contrasts and analyses of GLM1-A and GLM2-A to investigate whether the effects of sensory context and syllable categories can be explained away by modelling the effect of entropy.

At the 2^nd^ random effects level, we report results at *p_FWE_* < 0.05 cluster level corrected for the whole brain using an auxiliary voxel threshold of *p* < 0.001 uncorrected. For completeness we also present results using only a voxel threshold of *p* < 0.001(uncorrected) in the supplementary materials (**figure S4**).

## Results

### Behavioral results

#### Response distributions over the four choice options

Figure 2 shows the distribution of participants’ responses over the four choice options: ‘ba’, ‘da’, ‘ga’, ‘other’. An auditory ba stimulus is perceived mainly as a ‘ba’ percept, but on a fraction of trials also as a ‘da’ percept. Conversely, an auditory ga stimulus mainly evokes a ‘ga’ percept. Crucially, a ‘da’ percept is thus a possible perceptual interpretation for both auditory ba and visual ga stimuli (**figure 2A**). This perceptual uncertainty explains that participants can merge the conflicting McGurk signals (i.e., A_ba_ + V_ga_) into an illusory ‘da’ percept, because it is the perceptual interpretation that is possible for auditory and visual signals. By contrast, a visual ba is almost exclusively perceived as a ‘ba’ and an auditory ga as a ‘ga’ percept. As a result, participants cannot fuse these signals into a joint perceptual interpretation for both signals (**figure 2B**). Hence, on incongruent audiovisual trials (i.e., A_ga_ + V_ba_), participants segregate audiovisual signals and report the task-relevant ‘ga’ percept.

**Figure 2.**
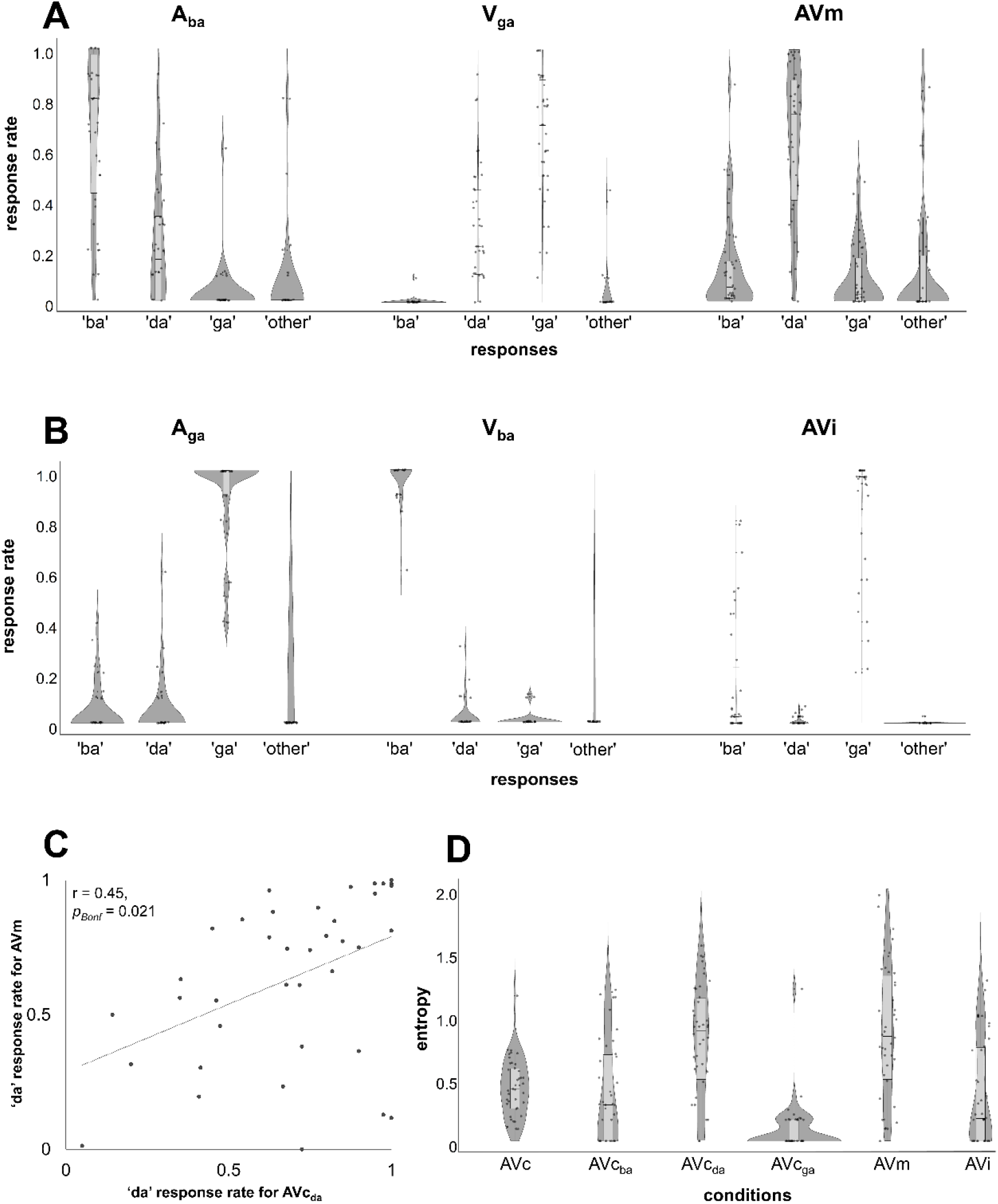
Behavioral results. (A) Response distribution for auditory ba, visual ga, and McGurk stimuli. (B) Response distribution for auditory ga, visual ba, and incongruent stimuli. (C) Correlation between the ‘da’ responses rate for McGurk stimulus (i.e., illusion rate) and ‘da’ responses rate for audiovisual congruent da stimuli (i.e., categorization accuracy). (D) Response entropy for audiovisual congruent syllables (pooled over ba, da, and ga stimuli), audiovisual congruent ba, audiovisual congruent da, audiovisual congruent ga, McGurk, and incongruent stimuli.

#### Response accuracy and illusion susceptibilities

A 3 (modality: A, V, and AVc) × 3 (syllable: ba, da, and ga) repeated measures ANOVA revealed significant main effects of modality (*F* = 42.25, *p* < 0.001, η² *=* 0.09) and syllable (*F* = 58.13, *p* < 0.001, η² = 0.31) and a significant interaction effect between modality and syllable (*F* = 18.45, *p* < 0.001, η² *=* 0.11) **(table S1)**. Overall, participants showed higher accuracy for the audiovisual congruent syllables than for the auditory or visual syllables. Likewise, they exhibited lower accuracy for da syllables than for ba and ga syllables. Post-hoc analysis showed that the categorization accuracy was higher for the ba syllable under AV congruent (AVc_ba_, mean = 0.90, SD = 0.12) than unisensory auditory presentation (mean = 0.67, SD = 0.32); the accuracy for the da syllable was higher for the audiovisual congruent (mean = 0.69, SD = 0.26) than the auditory (mean = 0.36, SD = 0.34) and visual modalities (mean = 0.43, SD = 0.27); the accuracy for ga syllable was higher for the AV congruent (mean = 0.98, SD = 0.05) than the visual condition (mean = 0.67, SD = 0.25).

The susceptibility to the McGurk illusion showed large interindividual variability, ranging from 0% to 100% ‘da’ percepts across observers. Moreover, observers’ McGurk illusion rate, i.e., their percentage of ‘da’ percepts on McGurk stimuli was significantly correlated with their percentage of veridical ‘da’ percepts for AVc_da_ stimuli (*Pearson’s r* = 0.45, *p_Bonf_* = 0.021, **figure 2C**). There was also a non-significant trend for a correlation with their percentage of veridical ‘da’ percepts for V_da_ stimuli (*Pearson’s r* = 0.37, *p_Bonf_* = 0.07).

#### Response entropy for unisensory and audiovisual stimuli

Figure 2D shows the response entropy for congruent stimuli (pooled over categories), for congruent stimuli separately for ba, da and ga syllables, McGurk, and incongruent audiovisual stimuli. While the overall entropy for congruent stimuli was lower than that for McGurk stimuli (t = - 5.66, *p* < 0.001, *Cohen’s d* = -0.87), there was no significant difference in entropy between the McGurk stimulus and the corresponding audiovisual congruent da stimulus (t = -0.79, *p* = 0.43, *Cohen’s d* = -0.12). This pattern suggests that observers successfully merge incongruent audiovisual McGurk signals into a unified ‘da’ percept whose perceptual uncertainty is comparable to that of their ‘da’ percept on congruent trials. It seamlessly aligns with previous findings that observers can be equally confident about their percept on McGurk and audiovisual congruent trials (Meijer and Noppeney, 2023).

The supplementary figure S2 comprehensively shows the response entropy separately for 3 (syllables: ba, da, ga) × 3(sensory modalities: A, V, AV) in addition to the McGurk and AV incongruent stimuli. A 3 (modality: A, V, and AVc) × 3 (syllable: ba, da, and ga) repeated measures ANOVA revealed significant main effects of modality (*F* = 20.19, *p*< 0.001, η² *=* 0.08) and syllable (*F* = 32.88, *p* < 0.001, η² = 0.15) and a significant interaction effect between modality and syllable (*F* = 30.35, *p* < 0.001, η² *=* 0.20) **(table S2)**. Follow-up t-tests for the simple main effects indicate a significant decrease in entropy for the audiovisual congruent ba relative to the auditory ba syllable (*t* = -4.36, *p_bonf_*< 0.001, *Cohen’s d = -*0.93), for the AV congruent da relative to the visual da syllable (*t* = -5.37, *p_bonf_*< 0.001, *Cohen’s d =* -1.15), and for the AV congruent ga relative to the visual ga syllable (*t* = -8.43, *p_bonf_* < 0.001, *Cohen’s d =* -1.80).

### fMRI results

#### Auditory and visual activations

Our results confirmed that auditory and visual stimuli increased activations along either the auditory (**GLM-1A, figure 3A - regions coded in red**) or visual (**GLM-1A, figure 3A - blue**) processing pathways that then converged in an extensive frontoparietal neural system shared across sensory modalities (**GLM-1A, figure 3A- pink**).

**Figure 3.**
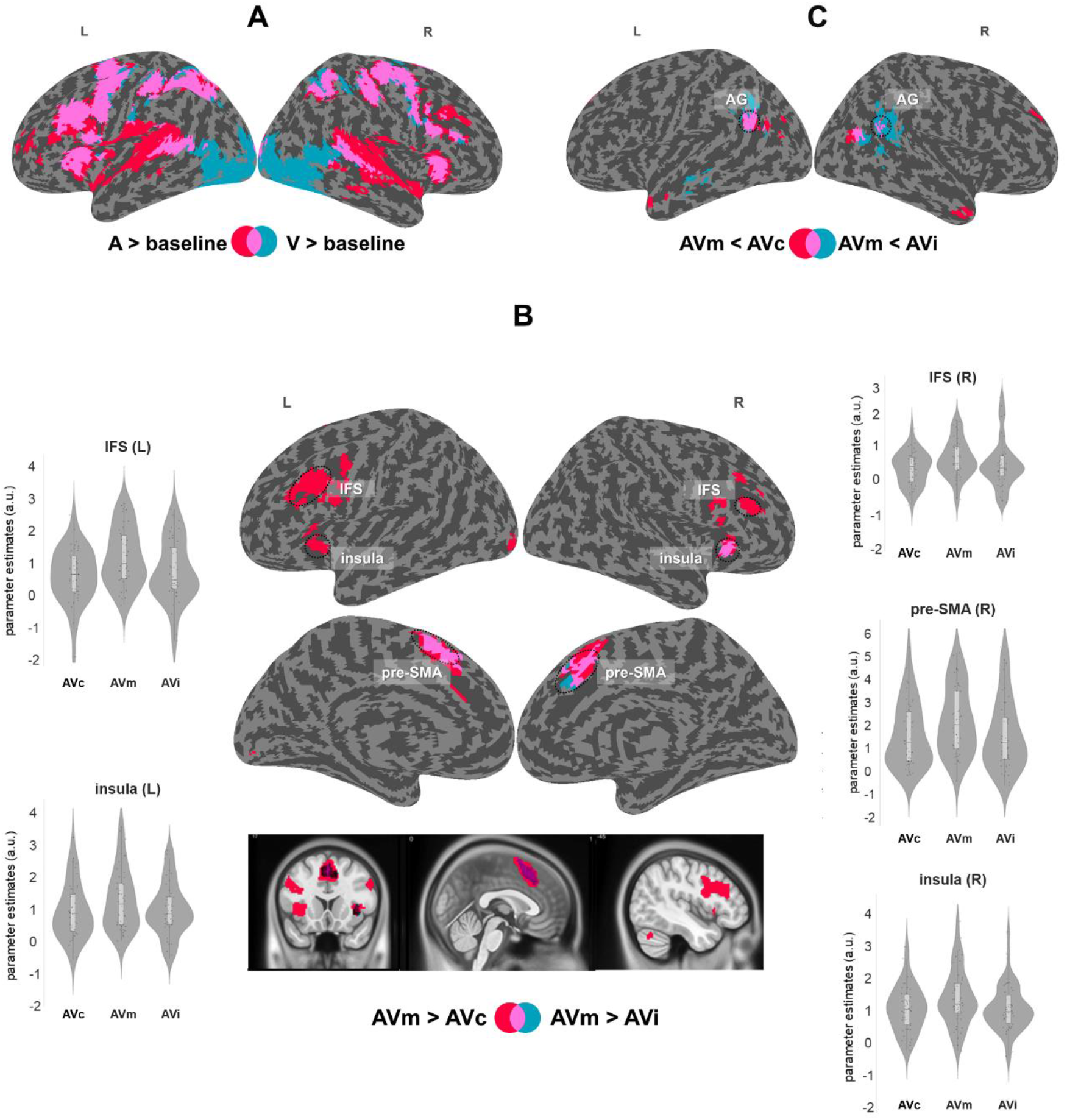
Effects of sensory contexts in GLM - 1A. (A) Increased activations for auditory (A, red) and visual (V, blue) stimuli relative to baseline, and their intersection (pink). (B) Middle: increased activations for McGurk stimuli (AVm) relative to audiovisual congruent (AVc, red) and incongruent (AVi, blue) stimuli, and their intersection (pink). left and right columns: violin plots showing the distribution over the subject-specific parameter estimates for AVm, AVc, and AVi relative to baseline at the MNI peak coordinate in left IFS (X = -46, Y= 26, Z = 28), right IFS (X = 38, Y= 36, Z = 20), right pre-SMA (X = 2, Y= 18, Z = 52), left insula (X = -40, Y= 16, Z = 0), and right insula (X = 30, Y= 24, Z = 6) defined by the AVm > AVc contrast. (C) Decreased activations for McGurk (AVm) relative to audiovisual congruent (AVc, red) and audiovisual incongruent (AVi, blue), and their intersection (pink). Activations are shown at *p*_FWE_ < 0.05 at the cluster level corrected for multiple comparisons within the entire brain, using an auxiliary uncorrected voxel threshold of *p* < 0.001. Thresholded images were rendered on an inflated canonical brain. Abbreviations: L - left, R - right, IFS - inferior frontal sulci, pre-SMA - pre-supplementary motor area, AG - angular gyri.

#### McGurk relative to audiovisual congruent stimuli

Compared to congruent stimuli, McGurk stimuli increased activations in IFS, pre-SMA extending into medial frontal gyrus, and insulae bilaterally, i.e., a network of regions involved in conflict processing and cognitive control (Duncan and Owen, 2000; Kerns et al., 2004; Brown and Braver, 2005). Further, they decreased activations in bilateral angular gyri and right posterior middle temporal gyrus **(GLM-1A, figure 3B, table - 1)**. In a follow-up analysis, we compared the BOLD-response for the McGurk stimulus that typically elicits an illusory ‘da’ percept selectively to the audiovisual congruent da stimulus that typically elicits a congruent ‘da’ percept. This more constrained comparison did not reveal any significant activations – thereby mirroring the behavioral results showing comparable response entropy for McGurk stimuli and audiovisual congruent ‘da’ stimuli.

**Table 1.**
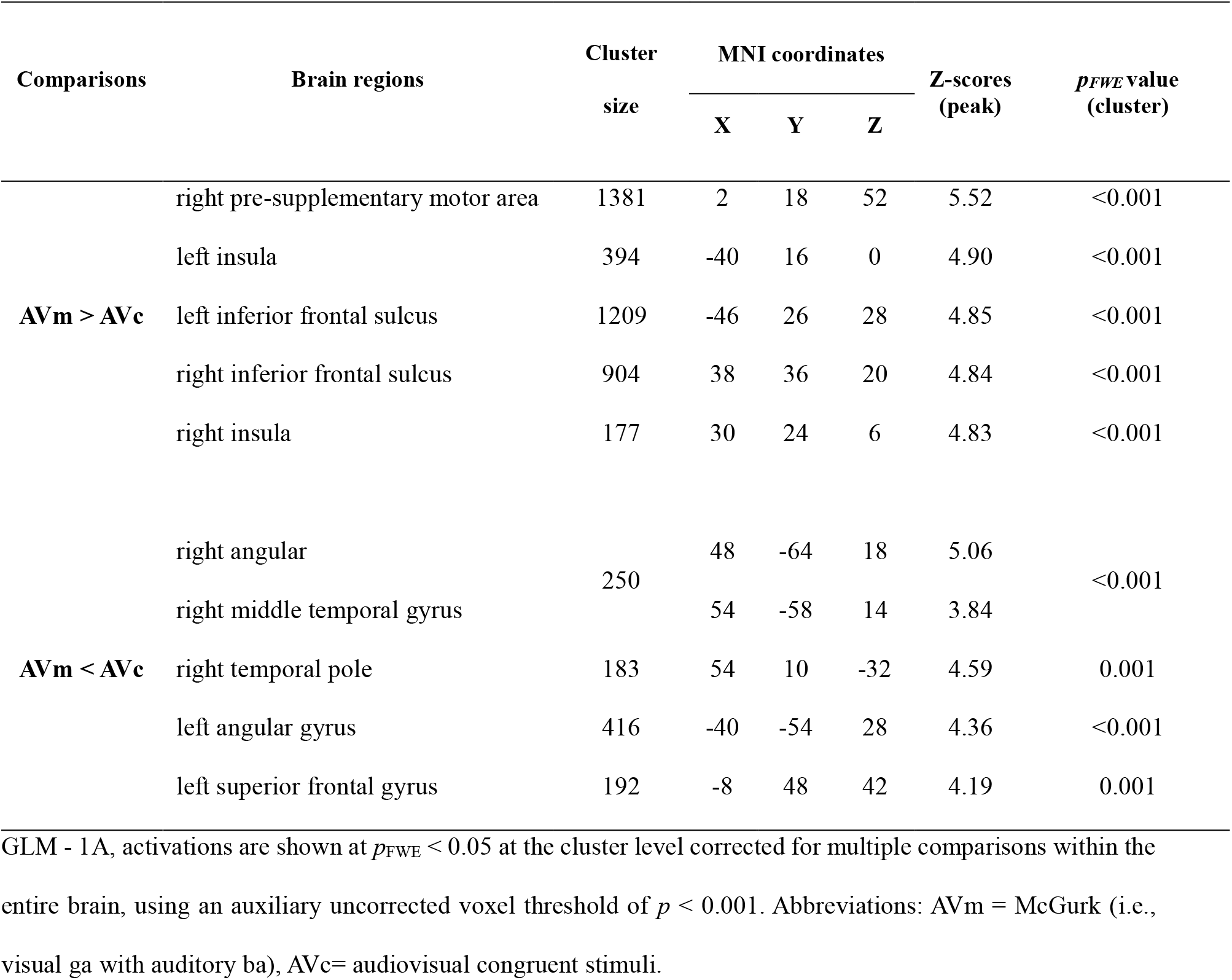
Brain activation differences between McGurk and congruent stimuli.

| Comparisons | Brain regions | Cluster<br>size | MNI coordinates |  |  | Z-scores<br>(peak) | *p*<sub>FWE</sub> value<br>(cluster) |
| --- | --- | --- | --- | --- | --- | --- | --- |
|  |  |  | X | Y | Z |  |  |
| **AVm > AVc** | right pre-supplementary motor area | 1381 | 2 | 18 | 52 | 5.52 | <0.001 |
|  | left insula | 394 | -40 | 16 | 0 | 4.90 | <0.001 |
|  | left inferior frontal sulcus | 1209 | -46 | 26 | 28 | 4.85 | <0.001 |
|  | right inferior frontal sulcus | 904 | 38 | 36 | 20 | 4.84 | <0.001 |
|  | right insula | 177 | 30 | 24 | 6 | 4.83 | <0.001 |
| **AVm < AVc** | right angular | 250 | 48 | -64 | 18 | 5.06 | <0.001 |
|  | right middle temporal gyrus |  | 54 | -58 | 14 | 3.84 |  |
|  | right temporal pole | 183 | 54 | 10 | -32 | 4.59 | 0.001 |
|  | left angular gyrus | 416 | -40 | -54 | 28 | 4.36 | <0.001 |
|  | left superior frontal gyrus | 192 | -8 | 48 | 42 | 4.19 | 0.001 |

GLM - 1A, activations are shown at $p_{FWE} < 0.05$ at the cluster level corrected for multiple comparisons within the entire brain, using an auxiliary uncorrected voxel threshold of $p < 0.001$ . Abbreviations: AVm = McGurk (i.e., visual ga with auditory ba), AVc= audiovisual congruent stimuli.

#### McGurk relative to audiovisual incongruent stimuli

Compared to incongruent stimuli, McGurk stimuli increased activations in pre-SMA extending into medial frontal gyrus and right insula and decreased activations in bilateral angular gyri and middle temporal gyri (**GLM-1A, figure 3C, table - 2**).

**Table 2.** Brain activation differences between McGurk and incongruent stimuli.

| Comparisons | Brain regions | Cluster<br>size | MNI coordinates | | | Z-score<br>s(peak) | $p_{FWE}$ value<br>(cluster) |
| --- | --- | --- | --- | --- | --- | --- | --- |
|  |  |  | X | Y | Z |  |  |
| AVm > AVi | left pre-supplementary motor area | 751 | -2 | 14 | 58 | 4.33 | <0.001 |
|  | right pre-supplementary motor area |  | 8 | 22 | 42 | 4.21 |  |
|  | right insula | 183 | 30 | 22 | -2 | 4.05 | 0.001 |
| AVm < AVi | right middle temporal gyrus | 1045 | 56 | -42 | 16 | 4.75 | <0.001 |
|  | right angular gyrus |  | 60 | -56 | 22 | 4.44 |  |
|  | left angular gyrus | 464 | -54 | -62 | 28 | 4.20 | <0.001 |
|  | left middle temporal gyrus | 106 | -56 | -18 | -14 | 3.97 | 0.022 |

the entire brain, using an auxiliary uncorrected voxel threshold of $p < 0.001$ . Abbreviations: AVm = McGurk
(i.e., visual ga with auditory ba), AVi = audiovisual incongruent (i.e. visual ba with auditory ga).

#### Activation differences across syllables

We observed activation differences across ba, ga and da syllables in widespread neural systems that were largely shared across auditory, visual and audiovisual stimuli. Key regions included the IFS/IFG, pre-SMA extending into medial frontal gyrus, and insulae bilaterally (for detailed results, see **table S3, Figure 4)**. Thus, activation differences across syllables arose in a network of regions that also exhibited differences between McGurk and congruent stimuli.

**Figure 4.**
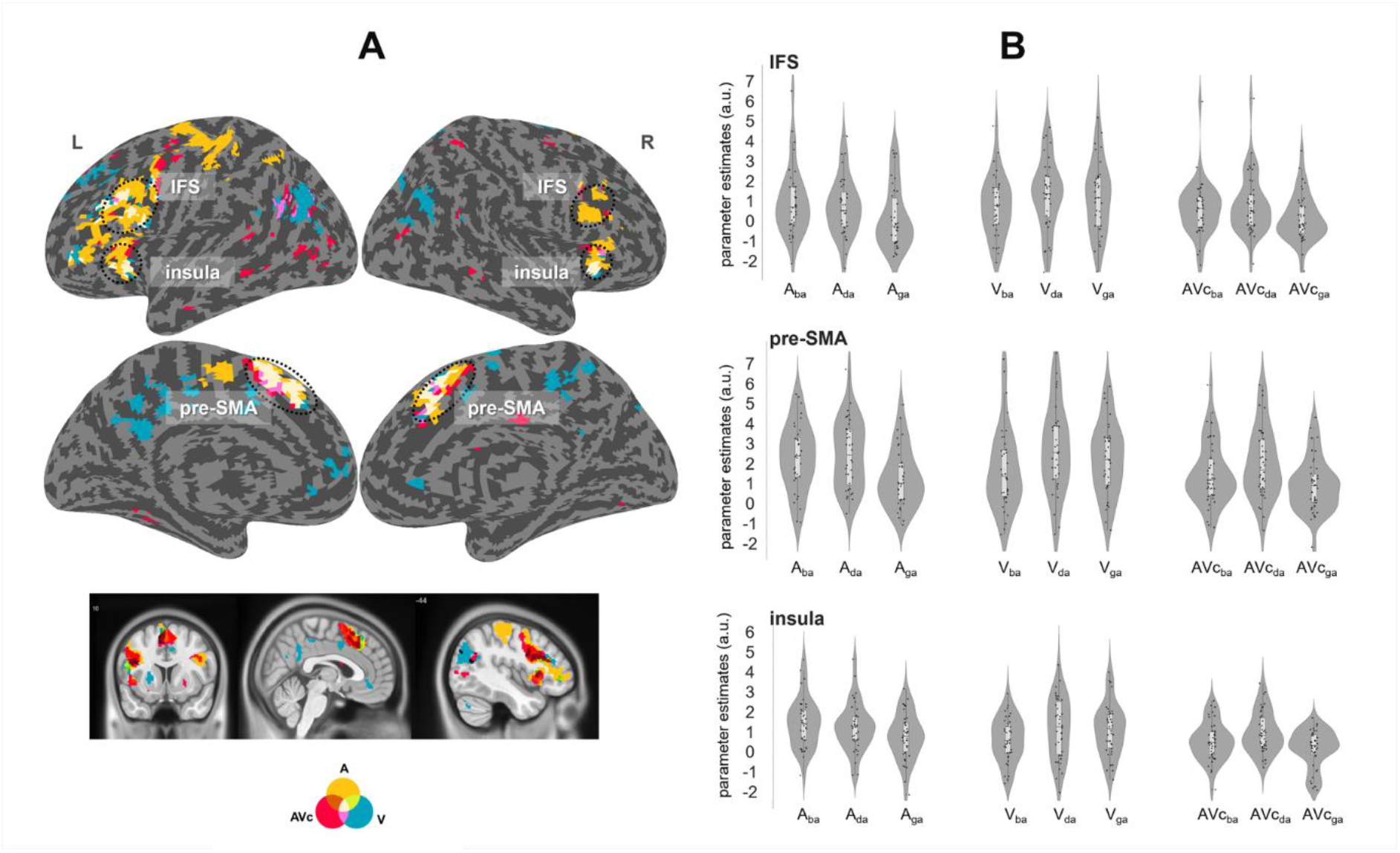
Effects of syllables in GLM - 2A. (A) Activation differences across syllables (ba, ga and da) separately for auditory (A, yellow), visual (V, blue), audiovisual congruent (AVc, red) stimuli, intersections between A and V (green), intersections between A and AV (wine dark red), intersections between V and AV (pink), and intersections between A, V, and AVc (white). (B) Parameter estimates for auditory, visual, and audiovisual congruent syllables relative to baseline at the MNI peak coordinate in left IFS (X = -46, Y= 26, Z = 28), right pre-SMA (X = 2, Y= 18, Z = 52), and left insula (X = -40, Y= 16, Z = 0) defined by the AVm > AVc contrast in GLM - 1A. Activations are shown at *p*_FWE_ < 0.05 at the cluster level corrected for multiple comparisons within the entire brain, using an auxiliary uncorrected voxel threshold of *p* < 0.001. Thresholded images were rendered on an inflated canonical brain. Abbreviations: L - left, R - right, IFS - inferior frontal sulci, pre-SMA - pre-supplementary motor area.

#### Effect of entropy over response distribution

Response entropy predicted activations in a widespread network of regions (GLM3, figure 5). The BOLD-response increased with greater entropy in bilateral IFS, pre-SMA extended to MFG, and insulae **(figure 5 – red, table 3)** and decreased in left angular gyrus and bilateral anterior MTG (**figure 5 - blue, table 3**).

**Figure 5.**
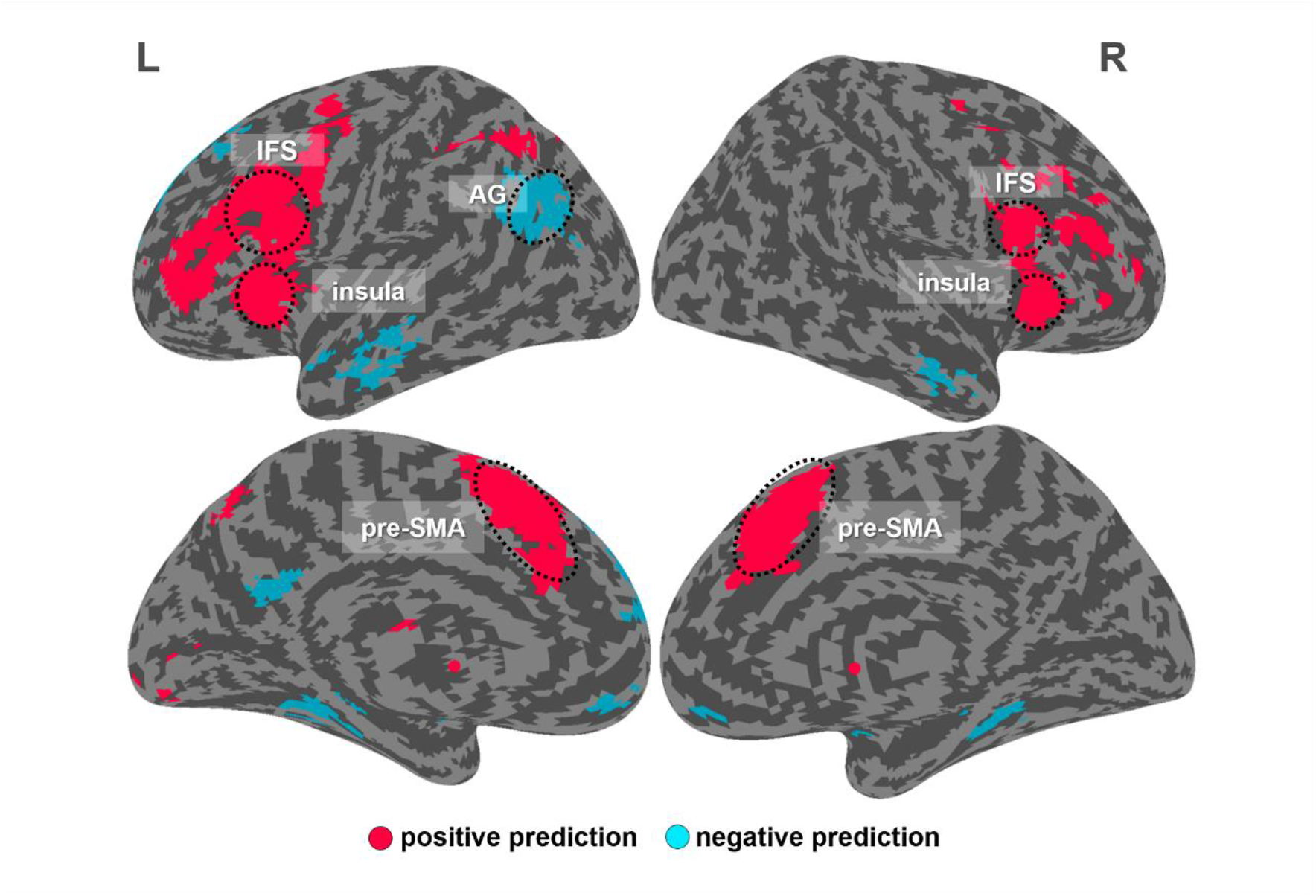
Effects of perceptual uncertainty (measured by entropy, GLM - 3). Increased activation with greater entropy over the response distribution (red), decreased activation with greater entropy over the response distribution (blue). Activations are shown at *p*_FWE_ < 0.05 at the cluster level corrected for multiple comparisons within the entire brain, using an auxiliary uncorrected voxel threshold of *p* < 0.001. Thresholded images were rendered on an inflated canonical brain. Abbreviations: IFS - left inferior frontal sulci, pre-SMA - pre-supplementary motor area, AG - angular gyrus.

**Table 3.** Brain activations predicted by entropy.

| Modulator | Brain regions | Cluster<br>size | MNI coordinates | | | Z-score<br>s(peak) | $p_{FWE}$ value<br>(cluster) |
| --- | --- | --- | --- | --- | --- | --- | --- |
|  |  |  | X | Y | Z |  |  |
| positive<br>prediction | right pre-supplementary motor area | 2260 | 4 | 18 | 50 | 7.32 | <0.001 |
|  | left insula | 5089 | -28 | 22 | 4 | 6.98 | <0.001 |
|  | left inferior frontal sulcus |  | -46 | 18 | 24 | 6.01 |  |
|  | right insula | 2244 | 32 | 24 | -6 | 6.63 | <0.001 |
|  | right inferior frontal sulcus |  | 46 | 8 | 24 | 5.90 | <0.001 |
| negative<br>prediction | left angular | 882 | -38 | -64 | 33 | 5.06 | <0.001 |
|  | left middle temporal gyrus | 463 | -58 | -16 | -26 | 4.63 | <0.001 |

GLM - 3, activations are shown at $p_{FWE} < 0.05$ at the cluster level corrected for multiple comparisons within the
entire brain, using an auxiliary uncorrected voxel threshold of $p < 0.001$ .

Further, we repeated the analyses and statistical comparisons described above (both whole brain and small volume corrected) using GLM-1B and GLM-2B that included entropy as one additional regressor. After accounting for variation in response entropy, none of the statistical comparisons revealed any significant results. This suggests that variation in response entropy across stimuli explains away activation differences across McGurk and congruent, incongruent stimuli as well as across syllables.

## Discussion

The McGurk illusion, a perceptual phenomenon that arises from merging conflicting audiovisual speech signals, has been a valuable tool for investigating how we understand speech (Sams et al., 1991; Sekiyama et al., 2003; Tuomainen et al., 2005; Tiippana et al., 2011; Tiippana, 2014; Alsius et al., 2018; Peelle, 2019). Some researchers however have recently argued that the neural and cognitive mechanisms underlying the McGurk illusion differ fundamentally from those involved in congruent audiovisual speech perception (Van Engen et al., 2017; Rosenblum, 2019; Getz and Toscano, 2021; Van Engen et al., 2022). Our study initially seems to support this view by showing activation increases for McGurk stimuli relative to congruent audiovisual stimuli in a network of areas typically associated with conflict monitoring and cognitive control processes (Duncan and Owen, 2000; Kerns et al., 2004; Brown and Braver, 2005). Intriguingly, these areas also exhibited greater activation for McGurk than incongruent trials, even though the latter are thought to place greater demands on conflict monitoring and control. Moreover, the level of activation also varied across ba, da and ga syllables, even when the auditory and visual signals were congruent. This finding suggests that even congruent speech can place varying executive demands. It aligns with our observation that perceptual uncertainty, measured by Shannon entropy over observers’ response distributions, strongly predicts the activation level in these areas, effectively explaining away differences between McGurk and normal congruent speech as well as differences between syllable classes.

Bayesian models view perception as inference based on noisy sensory signals (Helmholtz, 1867; Yuille and Bülthoff, 1993; Noppeney, 2021). Applied to speech perception, they propose that the brain infers a syllable from noisy acoustic signals and the corresponding facial movements (Magnotti and Beauchamp, 2017; Magnotti et al., 2018; Magnotti et al., 2020; Lindborg and Andersen, 2021; Kimmet et al., 2023; Meijer and Noppeney, 2023). Critically, the brain should integrate audiovisual signals from common sources, but segregate those from independent sources. Bayesian causal inference models deal with this so-called causal inference by computing estimates that fuse and segregate the signals. To account for observers’ uncertainty about the signals’ causal structure, they compute a final perceptual estimate by combining the fusion and segregation estimates weighted by the probabilities of common or independent sources. Bayesian causal inference models thereby intimately link observers’ causal and perceptual uncertainty (Kording et al., 2007; Shams and Beierholm, 2010; Noppeney and Lee, 2018; Noppeney, 2021).

In a syllable categorization task, perceptual and causal uncertainty arise from two distinct sources: i. the internal and external noise that corrupts the sensory signals and ii. the variability in how a particular phoneme or viseme is produced, (Bejjanki et al., 2011). This latter variability explains that the viseme ba is almost exclusively categorized as a ‘ba’, while the viseme ga is often confused with a ‘da’. Conversely, the auditory phoneme ga is almost always perceived as a ‘ga’, whereas the phoneme ba can also be perceived as a ‘da’ (**see figure 2A, B**). Hence, the McGurk illusion arises, because a ‘da’ percept is a possible interpretation for both the auditory ba and the visual ga signal. Further, this illusory ‘da’ percept is especially likely in participants that predominantly report a veridical ‘da’ percept for the audiovisual congruent da phoneme-viseme pairs (i.e., significant Pearson correlation). By contrast, the reverse pairing, such as a visual ba with an auditory ga in the incongruent trials, seldomly results in a fused ‘da’ percept, because such a ‘da’ interpretation is near-impossible for either auditory ga or visual ba inputs.

Crucially, understanding perception as inference based on noisy sensory signals dispenses with the categorical distinction between congruent, McGurk, and incongruent phoneme-viseme pairs. This perspective acknowledges that perceptual inference always carries a degree of uncertainty (Li and Ma, 2020). Even congruent phoneme-viseme pairs can generate seemingly conflicting sensory signals due to sensory noise and within-category variability. However, perceptual uncertainty is typically increased for McGurk stimuli because of their small non-noticeable intersensory conflict which invokes causal and perceptual uncertainty. Consistent with Bayesian principles, we indeed observed larger response entropies for McGurk than audiovisual congruent stimuli when pooled over all syllable categories. Perhaps surprisingly, perceptual uncertainty was also greater for McGurk than audiovisual incongruent conditions that introduce an even larger inter-sensory conflict. This difference most likely results from the fact that in the unisensory context the auditory ba syllable in McGurk stimuli is associated with a greater perceptual uncertainty than the ga syllable that is the auditory component in the incongruent stimuli.

In line with this entropy profile, we observed enhanced activations for McGurk relative to congruent trials in IFS, pre-SMA and insulae bilaterally - regions typically involved in conflict and cognitive control processes (Noppeney et al., 2008; Adam and Noppeney, 2010; Noppeney et al., 2010). Previous studies often attributed these activation increases to the intersensory conflict in McGurk stimuli and concluded that McGurk stimuli must therefore rely on distinct neural mechanisms (Wiersinga-Post et al., 2010; Perrachione and Ghosh, 2013). However, in our study activation differences were no longer significant when we compared McGurk stimuli to the congruent stimuli selectively from the corresponding da syllable. Further, activations in the same regions were also increased for McGurk relative to incongruent phoneme-viseme pairs, even though the latter should theoretically be associated with a greater perceived intersensory conflict. Activation levels in the same regions also varied across syllable categories of audiovisual congruent trials. Collectively, this pattern suggests that rather than being sensitive to inter-sensory conflict these regions respond more generically to perceptual uncertainty that differs not only across congruent, McGurk and incongruent stimuli, but also across syllable categories (**Figure S1, S2**). Indeed, in support of this conjecture, our follow-up analysis revealed a positive correlation between perceptual uncertainty (i.e., response entropy) and activation levels in exactly this network of regions (Figure 5). Moreover, including response entropy as a predictor in our regression models effectively explained away activation differences between McGurk and congruent/incongruent phoneme-viseme pairs.

Taken together, our results suggest that McGurk and congruent phoneme-viseme pairs rely on shared neural processing systems. As McGurk trials are often associated with greater perceptual uncertainty than congruent speech, they place greater demands on conflict and cognitive control processes and the associated network of regions. However, McGurk and audiovisual congruent da stimuli were associated with comparable response entropy and neural activity levels, even though the latter was physically congruent. These findings suggest that at least for our McGurk stimuli observers successfully fused the conflicting audiovisual signals into ‘da’ percepts that are comparable to their audiovisual congruent da counterparts. These behavioural and neuroimaging results dovetail nicely with a recent psychophysics and computational modelling study showing comparable perceptual and causal confidence for McGurk and congruent stimuli (Meijer and Noppeney, 2023).

In addition, the role of ‘classical multisensory areas’ such as superior temporal gyri/sulci (pSTG/S) in the processing of McGurk stimuli has been debated (Olson et al., 2002; Beauchamp et al., 2010; Erickson et al., 2014; Beauchamp, 2016; Hickok et al., 2018; Brang et al., 2020). Our data demonstrate that pSTG/S is sensitive to the incongruency of auditory and visual signals, exhibiting increased activation for the incongruent compared to the McGurk stimuli (**figure S3, table S4, GLM - 1A**). Similar results have previously been reported in studies focusing on linguistic (Jones and Callan, 2003; Bernstein et al., 2008; Szycik et al., 2009; Benoit et al., 2010; Nath and Beauchamp, 2012; Moris Fernandez et al., 2018; Murakami et al., 2018) and non-linguistic stimuli (Watson et al., 2013; Davies-Thompson et al., 2019). Specifically, one study using audiovisual emotional stimuli revealed that, while the increased pre-SMA/SMA activations were associated with task difficulty, the increased pSTG/S activations were related to audiovisual incongruency (Watson et al., 2013). Despite the substantial individual variation in the susceptibilities to the McGurk illusion, we did not find a significant correlation between the activation level of pSTS/G and observers’ illusion rate (Benoit et al., 2010; Nath et al., 2011; Nath and Beauchamp, 2012), which may be explained by differences in participants, stimuli (e.g., speaker), and tasks used in our experiment (Magnotti et al., 2015; Mallick et al., 2015; Alsius et al., 2018; Brown et al., 2018; Feng et al., 2019).

In conclusion, our research indicates that both the McGurk illusion and natural audiovisual speech perception result from inference based on noisy audiovisual signals. Consequently, both come with an inherent degree of uncertainty. Furthermore, both engage shared neural systems encompassing STS, IFS, pre-SMA and insulae. Notably, the McGurk stimuli increase activations in the latter IFS, pre-SMA and insular regions that form part of a wider cognitive control system. Critically, however, the activation differences between McGurk and congruent audiovisual stimuli within this cognitive control system can be directly attributed to observers’ degree of perceptual uncertainty. Collectively, our behavioral and neural results suggest that the McGurk illusion and natural speech perception lie on a continuous spectrum rather than being categorically different. From a practical viewpoint they support the validity of the McGurk illusion as a tool for studying natural audiovisual speech perception.

## Acknowledgments

This research was funded by the National Natural Science Foundation of China (No. 32171051). We thank Lizhen Qiu and Qi Yao for help with the data acquisition.

