## Supplementary material for "Perceptual uncertainty explains activation differences between audiovisual congruent speech and McGurk stimuli": Figure S1,Figure S2,Figure S3,Figure S4, Table S1, Table S2, Table S3, Table S4

#### Supplementary figures

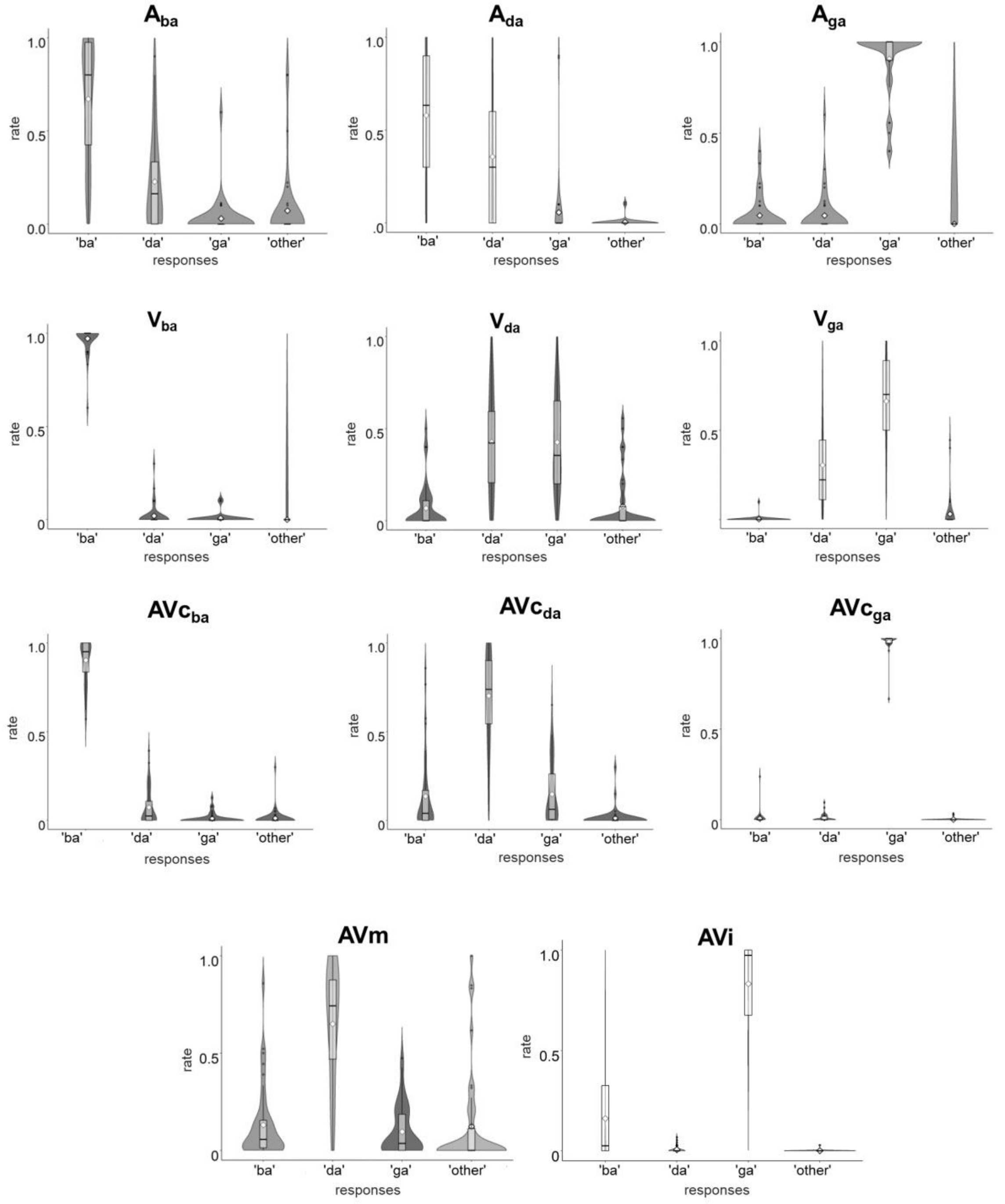


**Figure S1**. Distribution of responses over the four choice options ('ba', 'da', 'ga', and 'other') across the 11 stimulus classes. Abbreviations: $A_{ba}$ = auditory ba, $A_{da}$ = auditory da, $A_{ga}$ = auditory ga, $V_{ba}$ = visual ba, $V_{da}$ =

visual da, $V_{ga}$ = visual ga, $AVc_{ba}$ = audiovisual congruent ba, $AVc_{da}$ = audiovisual congruent da, $AVc_{ga}$ = audiovisual congruent ga, AVm = McGurk (i.e., visual ga with auditory ba), and AVi = audiovisual incongruent (i.e., visual ba with auditory ga).

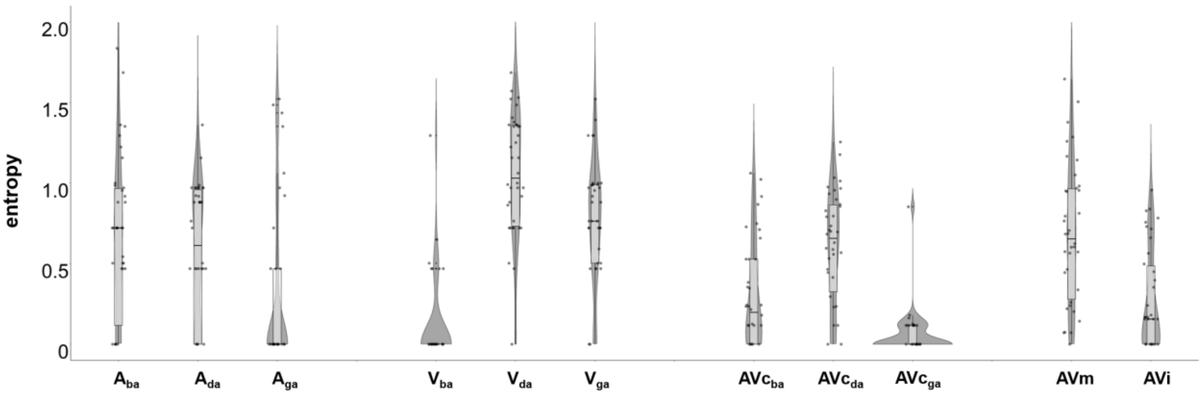


**Figure S2**. Response entropy for the 11 stimuli. Abbreviations: $A_{ba}$ = auditory ba, $A_{da}$ = auditory da, $A_{ga}$ = auditory ga, $V_{ba}$ = visual ba, $V_{da}$ = visual da, $V_{ga}$ = visual ga, $AVc_{ba}$ = audiovisual congruent ba, $AVc_{da}$ = audiovisual congruent da, $AVc_{ga}$ = audiovisual congruent ga, AVm = McGurk (i.e., visual ga with auditory ba), and AVi = audiovisual incongruent (i.e., visual ba with auditory ga)

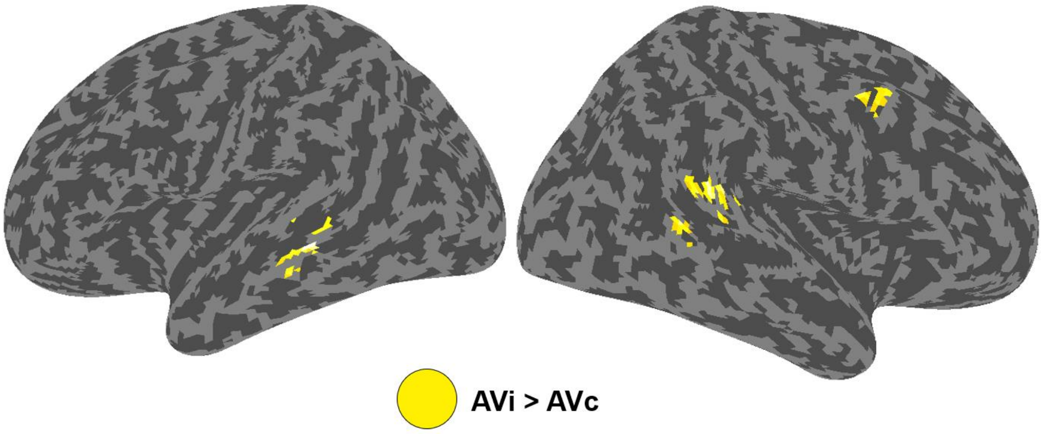


**Figure S3**. Increased activations for audiovisual incongruent (AVi) relative to audiovisual congruent stimuli (AVc, GLM - 1A). Activations are shown at $p_{FWE} < 0.05$ at the cluster level corrected for multiple comparisons within the entire brain, using an auxiliary uncorrected voxel threshold of $p < 0.001$.

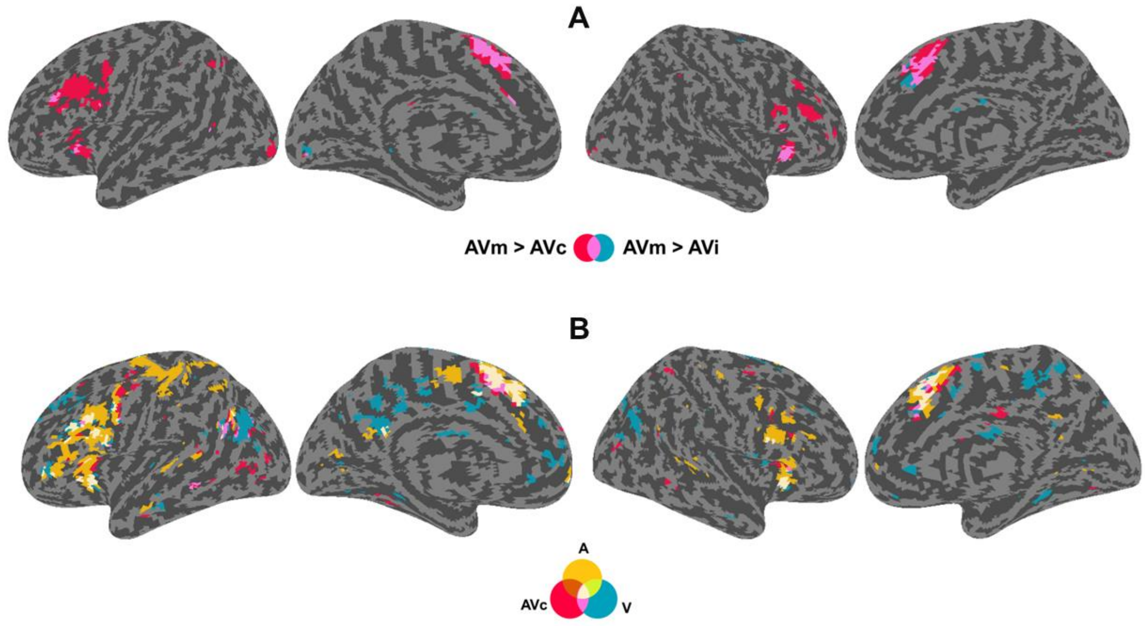


**Figure S4**. Effects of sensory contexts and phonemes at voxel threshold of $p < 0.001$ (uncorrected for illustrational purposes). (A) GLM - 1A: increased activations for McGurk stimulus (AVm) relative to audiovisual congruent (AVc, red) and incongruent (AVi, blue) stimuli, and their intersection (pink). (B) GLM - 2A: activation differences across syllables (ba, ga and da) for auditory (A, yellow), visual (V, blue), audiovisual congruent (AVc, red) stimuli, intersections between A and V (green), intersections between A and AV (wine dark red), intersections between V and AV (pink), and intersections between A, V, and AVc (white).

### Supplementary tables

**Table S1. Statistical results of the 3 (modality: A, V, and AVc) × 3 (syllable: ba, da, and ga) repeated measure ANOVA on categorization accuracy.**

| variables | F | $\eta^2$ | *p* | post-hoc comparisons | | t-value | Cohen's d | $p_{bonf}$ |
|---|---|---|---|---|---|---|---|---|
| **modality** | 42.25 | 0.09 | <0.001 | AVc | A | 8.93 | 0.92 | <0.001 |
| | | | | | V | 7.92 | 0.73 | < 0.001 |
| | | | | A | V | -1.57 | -0.19 | 0.371 |
| **syllable** | 58.13 | 0.31 | <0.001 | da | ba | -8.14 | -1.54 | < 0.001 |
| | | | | | ga | -9.01 | -1.56 | < 0.001 |
| | | | | ba | ga | -0.17 | -0.02 | 1 |
| **modality × syllable** | 18.45 | 0.11 | <0.001 | $AVc_{ba}$ | $A_{ba}$ | 4.81 | 0.98 | < 0.001 |
| | | | | | $V_{ba}$ | -1.56 | -0.322 | 1 |
| | | | | $AVc_{da}$ | $A_{da}$ | 7.09 | 1.45 | < 0.001 |
| | | | | | $V_{da}$ | 5.6 | 1.14 | < 0.001 |
| | | | | $AVc_{ga}$ | $A_{ga}$ | 1.6 | 0.33 | 1 |
| | | | | | $V_{ga}$ | 6.69 | 1.37 | < 0.001 |

Abbreviations: A = auditory stimuli, V = Visual stimuli, AVc = Audiovisual congruent stimuli, $A_{ba}$ = auditory ba, $A_{da}$ = auditory da, $A_{ga}$ = auditory ga, $V_{ba}$ = visual ba, $V_{da}$ = visual da, $V_{ga}$ = visual ga, $AVc_{ba}$ = audiovisual congruent ba, $AVc_{da}$ = audiovisual congruent da, and $AVc_{ga}$ = audiovisual congruent ga.

**Table S2. Statistical results of the 3 (modality: A, V, and AVc) × 3 (syllable: ba, da, and ga) repeated measure ANOVA on response entropy**

| variables | F | $\eta^2$ | *p* | post-hoc comparisons | | t-value | Cohen's d | $p_{bonf}$ |
|---|---|---|---|---|---|---|---|---|
| **modalities** | 20.19 | 0.08 | <0.001 | AVc | A | 3.74 | 0.49 | 0.001 |
| | | | | | V | 6.32 | 0.82 | < 0.001 |
| | | | | A | V | -2.58 | -0.34 | 0.035 |
| **syllable** | 32.88 | 0.15 | <0.001 | ba | da | -7.12 | -0.98 | < 0.001 |
| | | | | | ga | -0.19 | -0.03 | 1 |
| | | | | da | ga | 6.92 | 0.96 | < 0.001 |
| **modality × syllable** | 30.35 | 0.20 | <0.001 | $AVc_{ba}$ | $A_{ba}$ | -4.36 | -0.93 | < 0.001 |
| | | | | | $V_{ba}$ | 2.23 | 0.48 | 0.953 |
| | | | | $AVc_{da}$ | $A_{da}$ | 0.27 | 0.06 | 1 |
| | | | | | $V_{da}$ | -5.37 | -1.15 | < 0.001 |
| | | | | $AVc_{ga}$ | $A_{ga}$ | -2.75 | -0.59 | 0.234 |
| | | | | | $V_{ga}$ | -8.43 | -1.80 | < 0.001 |

Abbreviations: A = auditory stimuli, V = Visual stimuli, AVc = Audiovisual congruent stimuli, $A_{ba}$ = auditory ba, $A_{da}$ = auditory da, $A_{ga}$ = auditory ga, $V_{ba}$ = visual ba, $V_{da}$ = visual da, $V_{ga}$ = visual ga, $AVc_{ba}$ = audiovisual congruent ba, $AVc_{da}$ = audiovisual congruent da, and $AVc_{ga}$ = audiovisual congruent ga.

**Table S3. Brain activation differences across syllables separately for AV, A and V modalities**

| Comparisons | Brain regions | Cluster size | MNI coordinates X | Y | Z | Z-scores (peak) | $p_{FWE}$ value (cluster) |
|---|---|---|---|---|---|---|---|
| AV | left insula | 563 | -30 | 22 | 4 | 7.03 | <0.001 |
| | right pre-supplementary motor area | 1099 | -4 | 14 | 54 | 6.47 | <0.001 |
| | right insula | 212 | 34 | 24 | 4 | 6.20 | <0.001 |
| | left inferior frontal sulcus | 1024 | -42 | 10 | 24 | 5.85 | <0.001 |
| | right inferior frontal sulcus | 114 | 46 | 22 | 26 | 4.42 | 0.006 |
| | left angular gyrus | 192 | -46 | -60 | 26 | 3.72 | 0.001 |
| A | left insula | 2835 | -30 | 26 | -2 | 7.44 | <0.001 |
| | left inferior frontal sulcus | | -44 | 10 | 26 | 6.31 | <0.001 |
| | left pre-supplementary motor area | 1209 | -2 | 14 | 58 | 6.94 | <0.001 |
| | right insula | 491 | 30 | 22 | 0 | 5.96 | <0.001 |
| | right pre-supplementary motor area | 150 | 8 | 14 | 56 | 5.78 | <0.001 |
| | right inferior frontal sulcus | 567 | 44 | 10 | 32 | 5.52 | <0.001 |
| V | left pre-supplementary motor area | 925 | -2 | 22 | 48 | 6.22 | <0.001 |
| | right pre-supplementary motor area | | 6 | 24 | 42 | 4.93 | <0.001 |
| | left insula | 847 | -34 | 24 | -2 | 5.18 | <0.001 |
| | left inferior frontal sulcus | | -44 | 16 | 4 | 4.53 | <0.001 |
| | right insula | 174 | 30 | 24 | -4 | 5.12 | <0.001 |
| | left angular gyrus | 707 | -52 | -64 | 24 | 5.12 | <0.001 |
| | right angular gyrus | 426 | 56 | -64 | 28 | 4.44 | <0.001 |

GLM - 2A, activations are shown at $p_{FWE} < 0.05$ at the cluster level corrected for multiple comparisons within the entire brain, using an auxiliary uncorrected voxel threshold of $p < 0.001$. Abbreviations: A = auditory stimuli, V= visual stimuli, and AV = audiovisual congruent stimuli.

**Table S4. Brain activations for audiovisual incongruent > congruent stimuli**

| Comparisons | Brain regions | Cluster size | MNI coordinates X | Y | Z | Z-scores (peak) | $p_{FWE}$ value (cluster) |
|---|---|---|---|---|---|---|---|
| AVi > AVc | left middle temporal gyrus | 146 | -62 | -34 | 2 | 4.44 | <0.001 |
| | left superior temporal gyrus | | -50 | -40 | 2 | 3.54 | <0.001 |
| | right superior temporal gyrus | 202 | 60 | -40 | 16 | 4.17 | <0.001 |
| | right middle temporal gyrus | | 52 | -48 | 10 | 4.09 | <0.001 |
| | right middle frontal gyrus | 112 | 48 | 8 | 52 | 3.91 | <0.001 |

GLM - 1A, activations are shown at $p_{FWE} < 0.05$ at the cluster level corrected for multiple comparisons within the entire brain, using an auxiliary uncorrected voxel threshold of $p < 0.001$. Abbreviations: AVi = audiovisual incongruent stimulus, AVc= audiovisual congruent stimuli.
